# Origin and evolution of the Haustoriidae (Amphipoda): A eulogy for the Haustoriidira

**DOI:** 10.1101/2020.10.24.353664

**Authors:** Zachary B. Hancock, Hiroshi Ogawa, Jessica E. Light, Mary K. Wicksten

## Abstract

Haustoriid amphipods, despite their ubiquity in coastal sand or mud, have received little recent attention and their systematics and phylogenetics are largely unresolved. Some efforts have been made at classifying the family within the broader Amphipoda, but there is persistent incongruence in its placement among different authors and techniques. Furthermore, there exists no phylogenetic hypothesis of intrafamilial relationships despite the potential for rich biogeographic information to be gained given the specific habitat requirements of haustoriids and their limited dispersal abilities. In this work, we evaluate the competing hypotheses on the phylogenetic position of the Haustoriidae within Amphipoda by examining new and previously published sequences of nearly 100 species across 38 families. We find strong support for the Haustoriidae as basal gammarids, and that other families placed within the parvorder “Haustoriidira” are spread across Amphipoda. The radiation began during the Eocene and may have been driven in North America by the rapid filling of a coastal niche opened by the Chesapeake Bay impact crater. Unlike previous work, we find that the Pacific-endemic genus *Eohaustorius* is the most basal haustoriid, and that it separated from the rest of the family ~31 Mya. Finally, based on ancestral reconstructions, we provide taxonomic recommendations for relationships within Haustoriidae, including the elevation of a new genus, *Cryptohaustorius*. We conclude by recommending that the “Haustoriidira” be abandoned.

## INTRODUCTION

Amphipoda is an incredibly diverse order of crustaceans with nearly 10,000 described species (Barnard, 1957; Lowry & Myers, 2017). One such family of amphipods with uncertain affinity, the Haustoriidae, were deemed by J. Laurens Barnard as “perhaps the most interesting group of amphipods” (Barnard, 1969) likely because they are highly morphologically specialized to a fossorial lifestyle. Haustoriids are found on both open and protected beaches around the world but are hypothesized to have originated in the western North Atlantic (Bousfield, 1970). These amphipods have broad, fusiform bodies, lack eye pigmentation, and sport a dizzying array of spines and setae that give them the appearance of a fuzzy, opaque bean (**Fig. 1e**). They have lost article 4 on their mandibular palp (article 3 is strongly geniculate in compensation) and they are characterized by the absence of dactyls on pereopods 3–7. On gnathopods 1–2, the dactyls have been highly reduced; on gnathopod 1 they are a thin, curved nail, whereas they compose a minute chela on gnathopod 2. Pereopods 3–4 are perhaps the most curious: article 6 forms an expanded cup-like structure that is lined by stout spines for digging through the sediment (except in *Eohaustorius* spp.; pereopod 4 is a miniature version of pereopod 5). The last three legs are broadly expanded with each article as wide as they are long, and densely lined with comb setae. The posterior of haustoriids are as bushy as the front, and they have evolved powerful uropods with strong apical spines that aid in propelling them through the sand.

**Figure 1.**
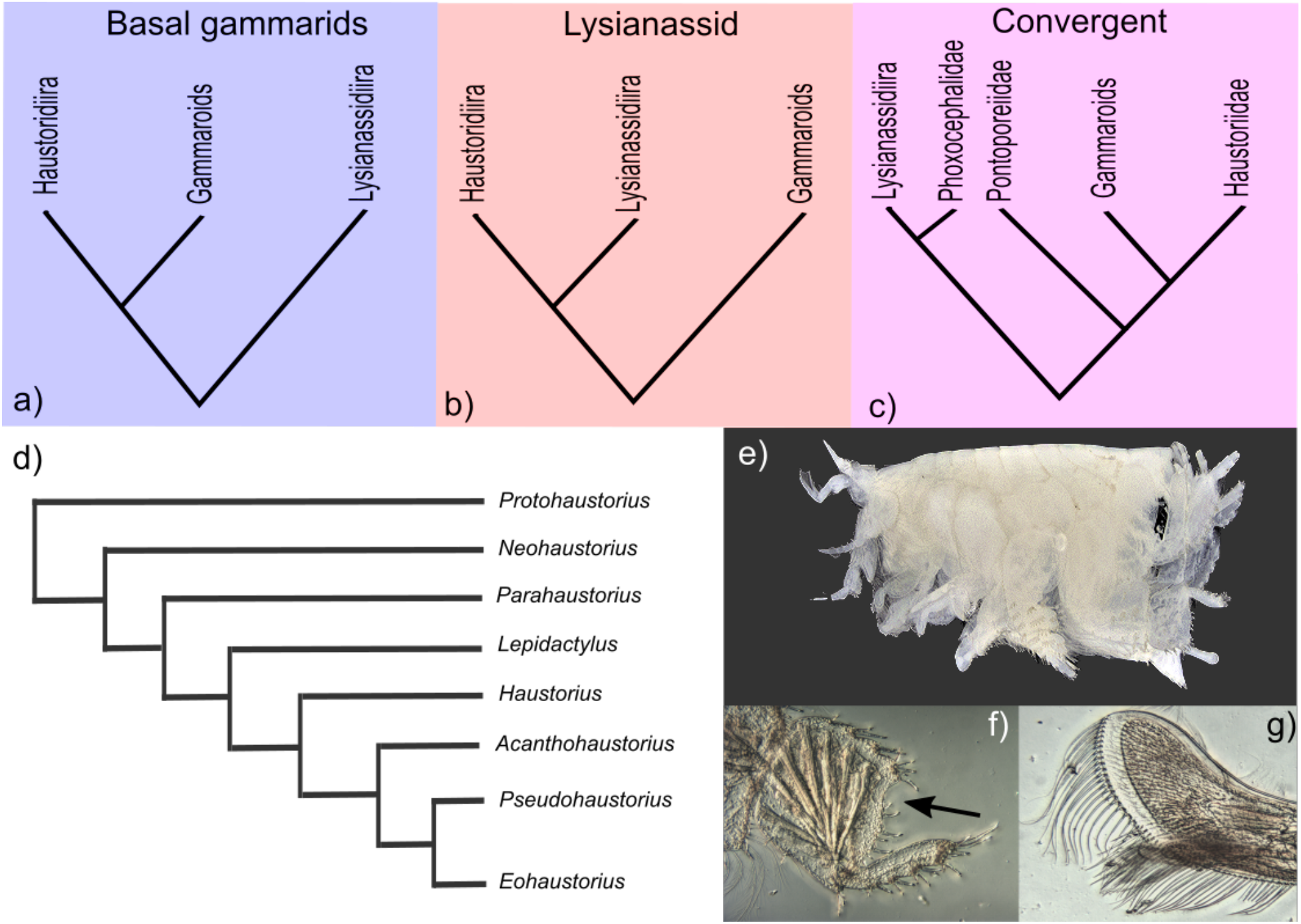
Phylogenetic hypotheses. a) basal gammarid hypothesis (Barnard & Drummond 1982); b) lysianassid hypothesis (Lowry & Myers 2017); c) convergent hypothesis; d) hypothesis of relationships within Haustoriidae (Sweet 1996); e) *Haustorius galvezi*; f) pereopod 6 article 5, arrow indicates taxonomically significant notch; g) maxilla 2 outer lobe compared to size of inner lobe (smaller, bottom).

Lowry & Myers (2013) erected a suborder of Amphipoda, Senticaudata, on the basis of the supposedly synapomorphic robust apical spines on uropods 1 and 2. This grouping includes many diverse and geographically widespread groups, such as the gammarids, talitrids, corophiids, and others. However, Lowry & Myers (2013) argue that the phoxocephalids and haustoriids have acquired this trait convergently and are excluded from this suborder. Instead, Lowry & Myers (2017) place the haustoriids in the suborder Amphilochidea, allying them with the pelagic lysianassids, the synopiids, and other fossorial amphipods included within the parvorder Haustoriidira (see *Taxonomy* in **Methods**). Verheye *et al*. (2016) has called into question the usefulness of the robust spines on the uropods for higher level classification, finding that it appears to evolve convergently across many families.

The modern taxonomy of the Haustoriidae (as with many amphipod families) was shaped by the morphological work of E.L. Bousfield and J. Laurens Barnard. Bousfield (1965) redefined the family from Stebbing’s (1906) original description, erecting two subfamilies: the Haustoriinae (or the “true” haustoriids) and the Pontoporiinae (previously their own family, the Pontoporiidae). In addition to the two haustoriid genera already recognized *(Haustorius* and *Lepidactylus*), Bousfield (1965) described five new genera of haustoriids and redistributed most of the described species at the time between them: *Protohaustorius, Acanthohaustorius, Pseudohaustorius, Neohaustorius*, and *Parahaustorius*. After 1965, the only species that remained in the nominal genus *Haustorius* were *H. arenarius*, the original type species, and *H. canadensis* Bousfield, 1962 (Hancock & Wicksten, 2018). These two would be joined later by *H. algeriensis* Mulot, 1967 from Algeria, *H. orientalis* Bellan-Santini, 2005 from the Mediterranean, *H. jayneae* Foster & LeCroy, 1991 from the eastern Gulf of Mexico, *H. mexicanus* Ortiz *et al*., 2001 from the state of Veracruz, Mexico, *H. galvezi* Hancock & Wicksten, 2018 from the western Gulf, and *H. allardi* Hancock & Wicksten, 2018 from the Mississippi Delta region. Among the haustoriids, the species within *Haustorius* are the largest by body size, and they tend to inhabit the high intertidal zone on surf-exposed sandy beaches (Hancock & Wicksten, 2018; Hancock *et al*., 2019). The proposed sister to *Haustorius* is *Lepidactylus*, which looks remarkably similar to *Haustorius* except for the lack of an overhang of epimeron 3 (Robertson & Shelton, 1980; Hancock & Wicksten, 2018). Hancock *et al*. (2019), using four molecular markers, found support for *Lepidactylus* as sister to *Haustorius*, but this work only included *L. triarticulatus* from the Gulf of Mexico and no other genera.

The phylogenetic position of the Haustoriidae within the Amphipoda has never been resolved. An early phylogenetic hypothesis was given by Barnard & Drummond (1982) in which they proposed that the Haustorioidea, a superfamily that includes the quintessential fossorial amphipod families (see *Taxonomy* in **Methods**), evolved from gammarid ancestors, and that the most basal lineage is the Pontoporeiidae (**Fig. 1a**). They wrote, “Pontoporeiids are very close to the Pontocaspian gammaroids… but differ in the enfeeblement of the gnathopods and the somewhat enlarged mandibular molars with relatively weaker triturative states and the loss of coxal gill 7” (Barnard & Drummond, 1982). Barnard & Drummond (1982) united the remaining haustorioids by the presence of the “haustoriid-like” pereopod 5: greatly expanded and heavily spinated. Those that did not then develop the “haustoriid-like” antenna were proposed to form a clade consisting of the Urothoidea, Platyischnopidae, and Condukiidae (Barnard & Drummond, 1982). Those that obtained the haustoriid antennae were then divided into two clades: 1) those that lack a mandibular palp and glassy spines (the Phoxocephalidae) and 2) those with: the Haustoriidae, Phoxocephalopsidae, Zobrachoidae, and the Urohaustoriidae.

A more recent phylogeny (Lowry & Myers, 2017), based again on morphological character states, proposed that the Haustoriidae are sister to the enigmatic Condukiidae, a family endemic to Australia, while retaining the basal position of the Pontoporeiidae (with the addition of an even more basal group, the Priscillinidae). However, Lowry & Myers (2017) found no support for a close relationship between the Zobrachoidae, Urohaustoriidae, Phoxocephalopsidae, and Haustoriidae; each of these families were scattered across the Haustorioidea tree.

A recent large-scale molecular phylogeny of Amphipoda comes from Copilaş-Ciocianu *et al*. (2020) in which they used four markers to infer the relationships of 210 species across 102 families. In their phylogeny, the only haustoriid included, *H. arenarius*, is found to be a basal gammarid with high support as suggested by Barnard & Drummond (1982) and in opposition to Lowry & Myers (2017). In addition, Copilaş-Ciocianu *et al*. (2020) concluded that the Haustoriidira is a polyphyletic parvorder consisting of multiple families spread across at least two different suborders (**Fig. 1c**). The lack of monophyly likely represents intense convergent evolution to a predominately shallow-water benthic lifestyle, a habitat shared by all of the included families. Several studies prior to Copilaş-Ciocianu *et al*. (2020) also found *H. arenarius* to be a basal gammarid (Verheye *et al*., 2016; Hou & Sket, 2017).

Lowry & Myers (2017) and Myers & Lowry (2018) have argued that molecular phylogenies that include some haustorioids tend to infer unprecedented relationships with little or no justification. For example, they note that *H. arenarius* is found to be the sister taxon to *Salentinella* (Bogidiellidae) in Verheye *et al*. (2016), a relationship that “During the three centuries that scientists have turned their attention to amphipod relationships, none of these associations have ever before been suggested” (Myers & Lowry, 2018). Further, no large phylogeny of Amphipoda to date based on molecular techniques has included more than a single haustoriid (*H. arenarius*), which may indicate that incomplete taxon sampling or long-branch attraction are impacting inferred topologies.

The relationships within Haustoriidae were evaluated by Sweet (1996) using morphological traits (**Fig. 1d**). He determined that *Protohaustorius* was the most basal haustoriid; a finding also suggested by Bousfield (1965). This genus most resembles the phoxocephalids and is easily diagnosable from the other haustoriids by its geniculate first antenna. The next group proposed to branch-off was *Neohaustorius* spp. followed by *Parahaustorius* spp. Sweet (1996) inferred that *Lepidactylus dysticus* was the outgroup to the remaining two clades of haustoriids: the *Haustorius* group and a hodgepodge group consisting of *Pseudohaustorius* spp., *Eohaustorius* spp. (a Pacific-endemic genus), and *Acanthohaustorius* spp. Sweet’s effort was the first and (prior to this work) only attempt at reconstructing a phylogeny of intrafamilial relationships of the Haustoriidae.

The first molecular phylogeny of the Haustoriidae focused on species endemic to the Gulf of Mexico (Hancock *et al*, 2019). This study found support for widespread cryptic diversity among haustoriid species, and a range of species delimitation techniques identified at least 4 independent lineages within the nominal species *L. triarticulatus*. Hancock *et al*. (2020) followed this work by expanding sampling to haustoriid species endemic to the North Atlantic and the Pacific. However, this work focused specifically on the evolution of genome size within the family and did not comment on the specific relationships inferred.

The bulk of haustoriid diversity is concentrated along the western North American coastline, which led Bousfield (1970) to postulate that the New England area was the center of origin for the family. Since then at least two dozen species have been described from the Gulf of Mexico (GoM) and Pacific coastlines, expanding the known distribution of haustoriids across the northern hemisphere. In light of this, haustoriids may not have been a rapid radiation along the North American coastline as Bousfield (1970) thought but may have begun to diversify long before when the continents were in closer proximity (i.e., the Cretaceous or earlier). Alternatively, the family itself could have a deeper origin with North American-specific genera appearing rapidly following the separation of the continents and isolation of the Atlantic from the Pacific Ocean, which had occurred by the Paleocene (56–61 Mya).

In this study, we examine three hypotheses concerning the phylogenetic affinity of Haustoriidae: 1) the Haustoriidira (haustoriids and their kin) are basal gammarids (Barnard & Drummond, 1982; **Fig. 1a**); 2) the Haustoriidira are a sister group to the lysianassids (Lowry & Myers, 2017; **Fig. 1b**); and 3) that the Haustoriidira is polyphyletic, with families spread across the Amphipoda (Verheye *et al*., 2016; Hou & Sket, 2017; Copilaş-Ciocianu *et al*., 2020; **Fig. 1c**). Finally, we provide taxonomic recommendations to aid in resolving incongruences between molecular and morphological phylogenies of haustoriid amphipods.

## MATERIALS AND METHODS

### TAXONOMY

The current taxonomic hypothesis of the Haustoriidae is based on Lowry & Myers (2017) and is as follows:

**Order** Amphipoda Latreille, 1816
**Suborder** Amphilocidea Boeck 1871
**Infraorder** Lysianassida Dana, 1849
**Parvorder** Haustoriidira Stebbing, 1906
**Superfamily** Haustorioidea Stebbing, 1906

**Family** Cheidae Thurston, 1982
**Family** Condukiidae Barnard & Drummond, 1982
**Family** Haustoriidae Stebbing, 1906

**Genus** *Acanthohaustorius* Bousfield, 1965*
**Genus** *Eohaustorius* Barnard, 1957*
**Genus** *Haustorius* Müller, 1775*
**Genus** *Lepidactylus* Say, 1818*
**Genus** *Neohaustorius* Bousfield, 1965*
**Genus** *Parahaustorius* Bousfield, 1965*
**Genus** *Protohaustorius* Bousfield, 1965*
**Genus** *Pseudohaustorius* Bousfield, 1965*
**Family** Ipanemidae Barnard & Thomas, 1988
**Family** Otagiidae Hughes & Lörz, 2013
**Family** Phoxocephalidae Sars, 1891*
**Family** Phoxocephalopsidae Barnard & Drummond, 1982
**Family** Platyischnopidae Barnard & Drummond, 1979
**Family** Pontoporeiidae Dana, 1852*
**Family** Priscillinidae d’Udekem d’Acoz, 2006
**Family** Sinurothoidae Ren, 1999
**Family** Urohaustoriidae Barnard & Drummond, 1982
**Family** Urothoidae Bousfield, 1978*
**Family** Zobrachoidae Barnard & Drummond, 1982

All designations marked with “*” indicates that a member of the family or genus was included in this study.

### SAMPLE COLLECTION

Samples from the United States were collected using methods in Hancock *et al*. (2020) and were preserved in 95% ethanol. *Eohaustorius estuarius* from the Pacific coast was collected by Gary Buhler of Northwest Amphipod, LLC. Specimens from Japan were collected from three sites: 1) Nakusa-no-hama beach, Kemi, Wakayama City, Wakayama Prefecture; 2) Ustumi, Minamichita town, Chita district, Aichi Prefecture; and 3) Banzu tidal flat, located at the mouth of Obitsu River, Chiba Prefecture. Amphipods were taken together with the sandy mud substrate of the tidal flat surface by a shovel at low tide in the intertidal zone or were caught by a trawl net (net length: 10,000 mm; mesh size: 1 mm; width of mouth flame: 500 mm; height of mouth flame: 250 mm, with front chain; products by RIGO Co., Ltd.) at high tide in the subtidal zone. Samples were sieved through a 1mm mesh and sorted in the field, then fixed and preserved in 70–99% ethanol. *Eohaustorius* sp. and additional samples of *E. subulicola* were collected by Aoi Tsuyuki of Hokkaido University.

### DNA EXTRACTION AND DATASETS

Whole genomic DNA was extracted from either the whole specimen or pereopods 5–7 using an EZNA Tissue Kit (Omega Bio-tek Inc.) following the manufacturer’s protocols. Mitochondrial cytochrome oxidase I (*COI*) and nuclear 28S ribosomal RNA (*28S*) and histone H3 (*H3*) were amplified via polymerase chain reaction (PCR). Primer sets and PCR conditions for *COI* and *28S* follow Hancock *et al*. (2019); conditions for *H3* follow Esmaeili-Rineh *et al*. (2015). Amplicons were verified using gel electrophoresis and we used ExoSAP-IT (Affymetric Inc.) to purify positive PCR products. Sanger sequencing on forward and reverse strands was performed at DNA Analysis Facility on Science Hill at Yale University. Sequences were manually edited in Sequencher v.4.10.1 (Gene Codes Corp.).

In addition to samples collected specifically for this study, we incorporated datasets from three studies: Hancock *et al*. (2019), Hancock *et al*. (2020), and Copilaş-Ciocianu *et al*. (2020). We subset the latter dataset (originally with 210 species) to include only the clades designated “Gammaroids”, “Miscellaneous”, “Lysianassoids”, and “Crangonyctoids”, the last of which acted as an outgroup (see Figure 1 in Copilaş-Ciocianu *et al*, 2020). Combined, these studies include 95 species across 38 families, including ~50% of recognized haustoriid species and at least one species for each described genus in the family (**Table 1**). We performed alignments using MAFFT v7 (Katoh & Standley, 2013) and checked *COI*and *H3* in Mesquite 3.5 (Maddison & Maddison, 2018) for the presence of premature stop codons. We found evidence of saturation at the third codon position in *COI* (**Fig. S1**); this position was removed from subsequent analyses. The final alignment length of *COI* was 458 bp; *H3* was 376 bp; and *28S* was 1,799 bp; total alignment length of 2,633 bp.

**Table 1.**
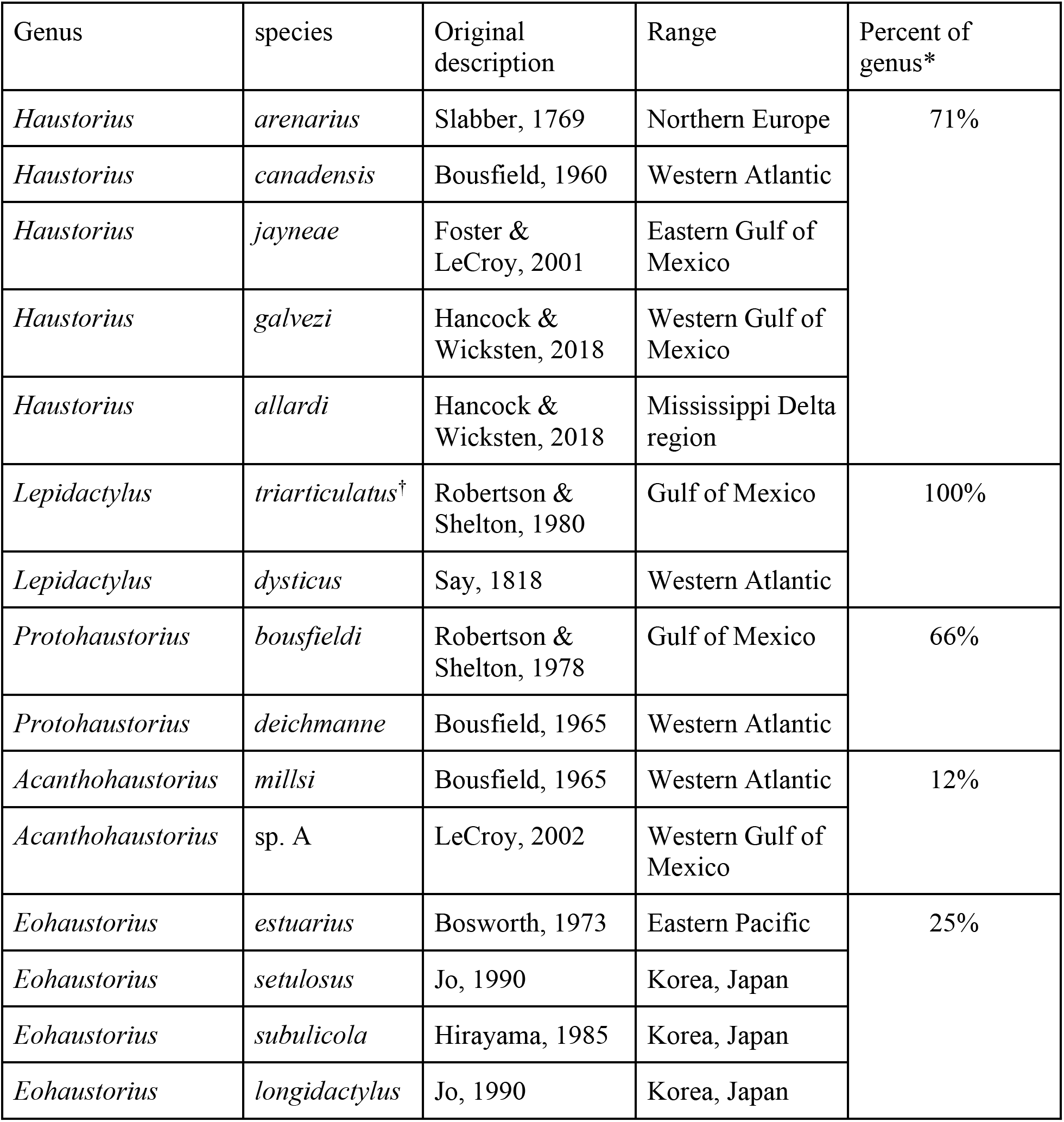
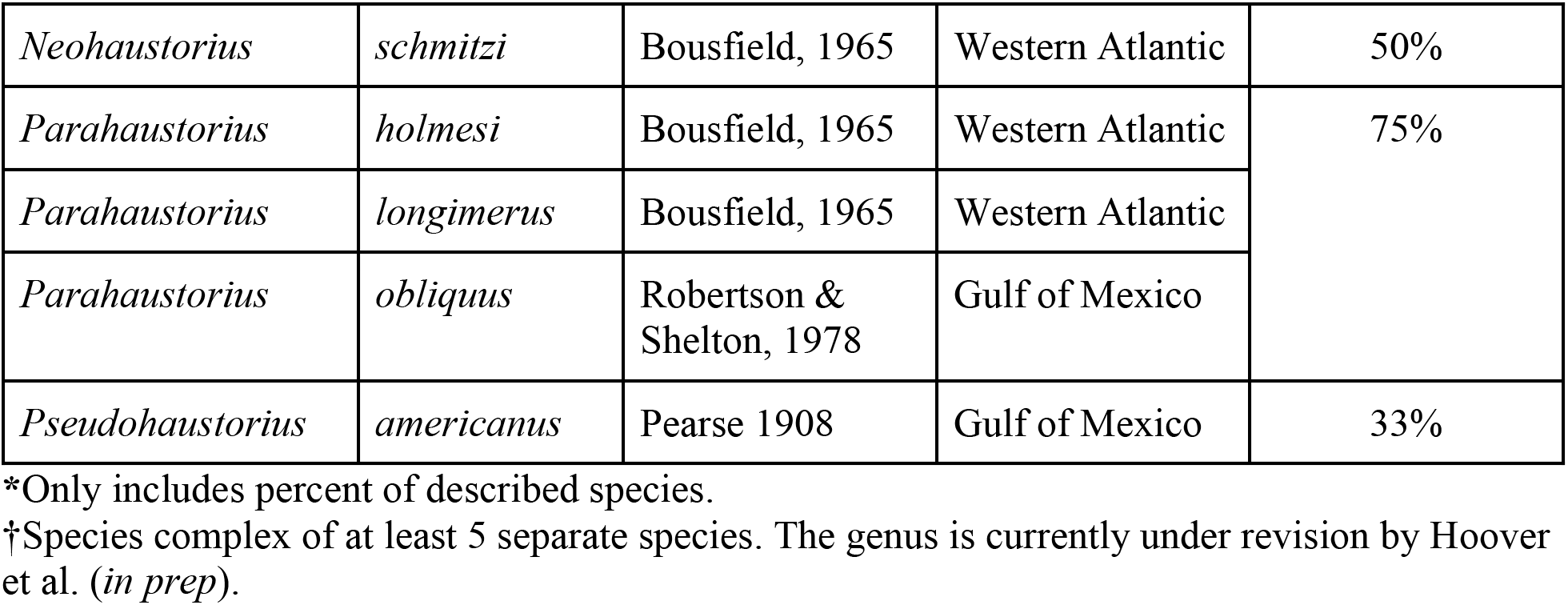
Haustoriid species included in this study.

To analyze relationships within the Haustoriidae, we included the loci above and two additional loci, *18S* and mitochondrial ribosomal *16S*. These sequences came from Hancock *et al*. (2019), and we reduced the datasets to include only one individual per species.

### PHYLOGENETIC METHODS

Phylogenetic inference was performed using two maximum-likelihood (ML) methods, IQ-TREE v2.1.3 (Minh *et al*., 2020) and RAxML v.8 (Stamatakis, 2014), and a Bayesian method, MrBayes (Huelsenbeck & Ronquist, 2001). Phylogenetic analyses were performed on the full concatenated dataset (*n* = 95) in which each gene was a separate partition. For the protein-coding genes, we designated a GTR + I + G model of substitution; an HKY model was set for *28S*. These models were identified as the best model by PartitionFinder2 (Lanfear *et al*., 2017). To assess node support in IQ-TREE, we performed 1,000 ultrafast bootstrap replicates (UB) as well as the Shimodaira-Hasegawa approximate likelihood ratio test (SH-aLRT; Shimodaira & Hasegawa, 1999). In RAxML, we performed rapid bootstrapping under the GTRCAT model with 1,000 replicates. For our Bayesian analysis in MrBayes, we designated a Birth-Death tree prior and ran two independent MCMC chains of 10 million generations each with a 25% burn-in to ensure ESS values > 200. Consensus trees were generated using the command *sumtrees* in DendroPy (Sukumaran & Holder, 2010).

A large number of sequences failed the composition *χ*^1^ test (41 sequences, ~43%) implemented in IQ-TREE, and 50 sequences (~52%) had missing data >50%. This was due to a combination of large gaps in some of the aligned *28S* sequences and some samples lacking a *28S* sequence altogether. Thus, we performed the above methods on a reduced dataset that only contained those that passed the composition *χ*^1^ test (*n* = 54) to evaluate how missing data may be impacting our results.

To investigate relationships within the Haustoriidae, we performed species-tree inference using BPP (Rannala & Yang, 2003). A guide-tree was produced using IQ-TREE on the reduced haustoriid-only dataset. Tree search was performed using subtree pruning and regrafting (SPR), and the MCMC was run for 200,000 generations sampling every 2 generations with a 10,000 generation burn-in. We then generated a maximum-clade credibility tree from the set of trees produced during the MCMC using the command *sumtrees* in DendroPy.

Finally, we also inferred a cladogram using a modified character matrix of 29 traits from Sweet (1996) of the Haustoriidae. We altered the original matrix by removing traits specific to the outgroup and any apomorphic trait. We additionally modified some trait scores to align with identified taxonomically important traits from Hancock & Wicksten (2018). The cladogram was inferred by a heuristic search of the 100 most parsimonious trees in Mesquite. Tree-space was searched using SPR. A majority-rule consensus tree was generated from the 100 trees.

### DIVERGENCE-TIME DATING

To estimate the age of the Haustoriidae, we used BEAST2 on the full concatenated dataset (*n* = 95) and applied 5 calibration points from Copilaş-Ciocianu *et al*. (2020) and Hancock *et al*. 2020. Age ranges to follow represent the 95% highest posterior density (HPD). We first set an exponential prior on the age of the Ponto-Caspian gammarids as 9–83 million years ago (Mya), corresponding to amber fossils in the Caucasus (Derzhavin, 1927; Copilaş-Ciocianu *et al*. 2020). Next, following Copilaş-Ciocianu *et al*. (2020), we applied exponential priors to the age of the Crangonyctidae-Pseudocrangonyctidae and the Niphargidae-Pseudoniphargidae splits according to Baltic amber fossils from the Eocene as 38–215 Mya. We then set exponential priors on splits within the Haustoriidae based on Hancock *et al*. 2020: 1) the proposed closure of the Okefenokee Trough ~1.75 Mya (set as 1.55–4.5 Mya; Bert, 1986; Avise, 1992; Knowlton & Weigt, 1998); and 2) the hypothesized Pleistocene colonization of Europe by *H. arenarius* (0.26–15 Mya; Bousfield, 1970).

### ANCESTRAL HABITAT AND MORPHOLOGY RECONSTRUCTION

To identify taxonomically informative character traits informed by our molecular phylogenies, we reconstructed ancestral character traits in R using the package *phytools* (Revell, 2012). We focus on the following traits that have emerged in the literature as being informative at the genus level: 1) epimeron 3 overhangs the urosome (1) or not (0); 2) maxilla 2 outer plate the same size (0), slightly larger (1), or twice as large as the inner (2); 3) presence (1) or absence (0) of a lobe on article 5 of pereopod 6; 4) number of spine groups on the posterior margin of article 4 on pereopod 7: 1, 2, 3, or 4; and 5) telson is uncleft (0), marginally cleft (1), or cleft to the base (2).

We also inferred the ancestral habitat type by coding each species according to whether they occurred on the open coast (“open”) or in brackish estuaries (“bay”). Furthermore, we searched for phylogenetic trends related to bottom preference: 1) fine sand; 2) medium/coarse sand; and 3) mud. Finally, we reconstructed the ancestral state of whether species occurred in the intertidal zone or subtidally. The complete table with character states, habitat information, and sample location can be found in the supplementary materials (**Table S1**).

## RESULTS

### PHYLOGENETIC RESULTS

All analyses of the phylogenetic position of the Haustoriidae within Amphipoda found strong support for a position sister to the gammaroids with the lysianassids as an outgroup (**Fig. 2**). We found that this result was not influenced by missing data as the reduced analysis also found this relationship (with even stronger support, **Fig. S2–5**). Furthermore, we found that the other families assigned to Haustoriidira that were included in our analyses (Pontoporeiidae, Phoxocephalidae, and Urothoidae) showed no affinity to the true haustoriids (**Fig. 2**). Instead, these families were found to be basal lysianssids (Pontoporeiidae, along with Bathyporeiidae) or belonged to clades independent of both the lysianassids and the gammaroids (i.e., Phoxocephalidae and Urothoidae).

**Figure 2.**
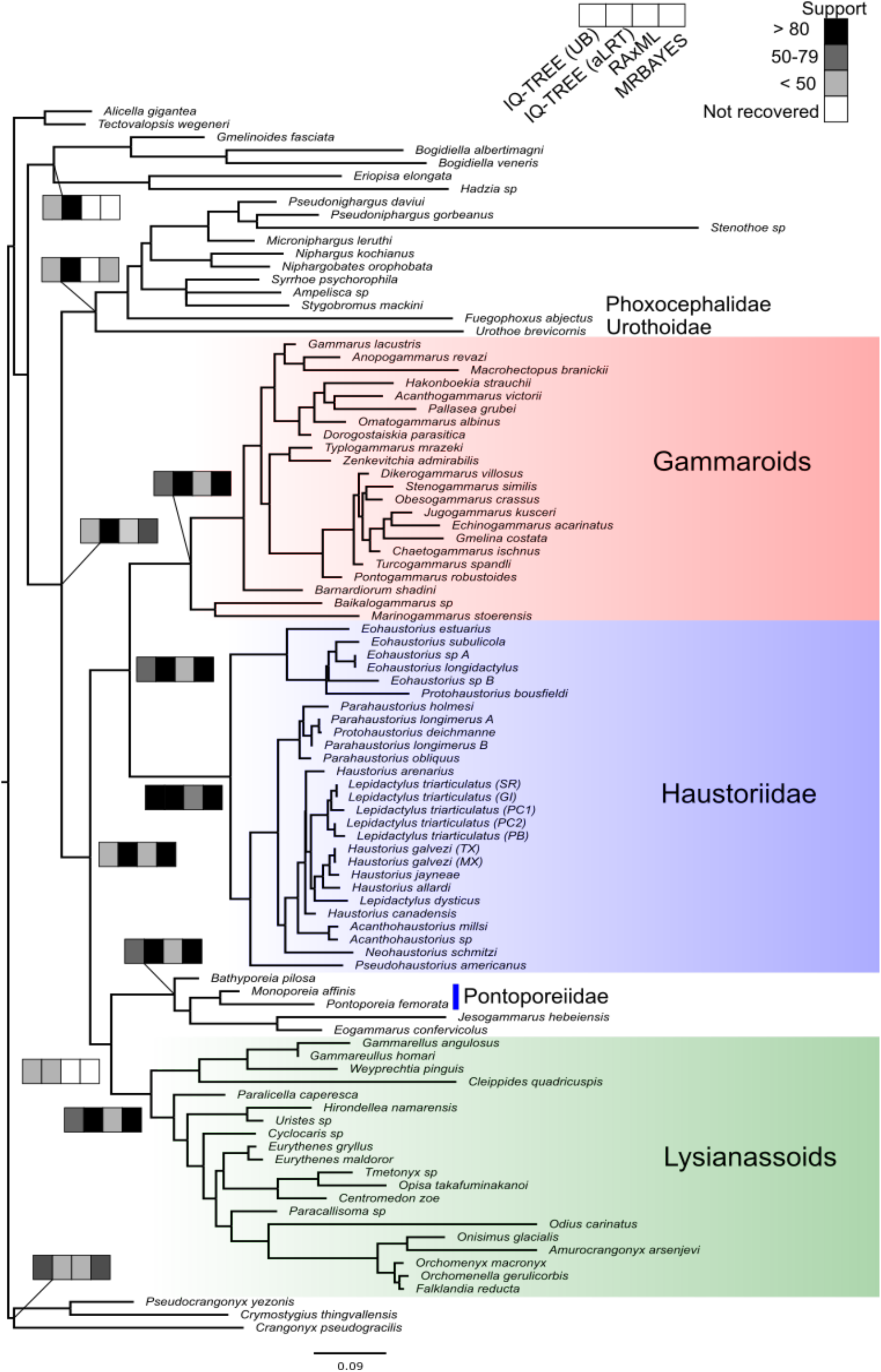
Maximum-likelihood phylogeny of Amphipoda from IQ-TREE.

The BPP analysis found that the type-genus *Haustorius* is paraphyletic with respect to *Lepidactylus* with high support (PP = 1.0), with *L. dysticus* grouping with the *Haustorius* species endemic to the GoM. The GoM endemic *Lepidactylus* – the *L. triarticulatus* species complex – was monophyletic, but distantly related to the *L. dysticus* + GoM endemic *Haustorius* clade (**Fig. 3**). The remaining *Haustorius* species, *H. canadensis* and *H. arenarius*, were found to be sister taxa and an outgroup to the aforementioned clades. The relationships between this “*Haustorius-Lepidactylus*” clade and the remaining genera remains uncertain due to relatively low node support (PP < 0.6). However, we did recover the monophyly of the remaining genera (except for *Protohaustorius*) and found strong support for the Pacific-endemic genus, *Eohaustorius*, as the earliest split from the other haustoriids.

**Figure 3.**
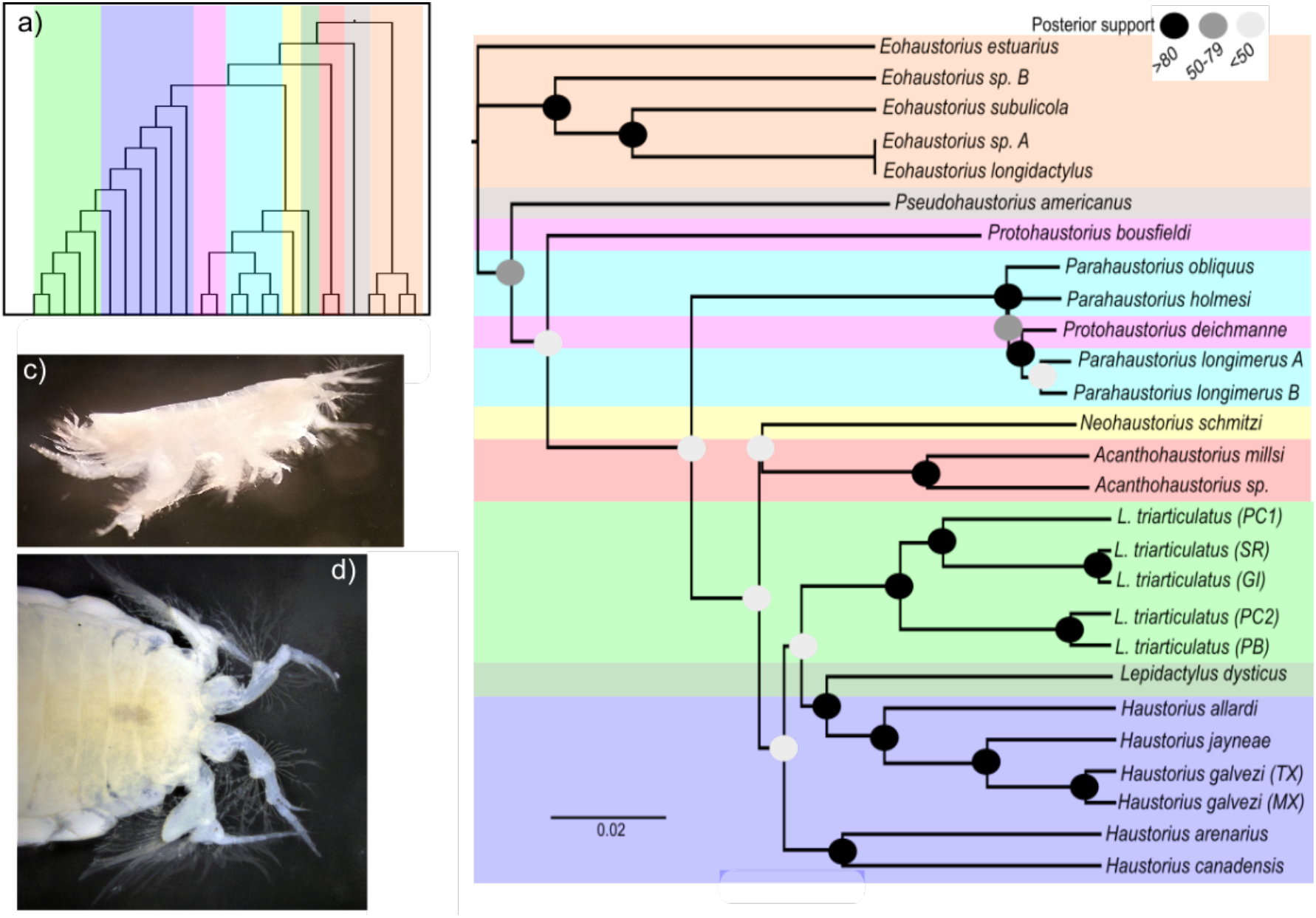
Species tree of Haustoriidae from BPP. Colors represent nominal genera. a) cladogram from the character matrix, colored clades match those from the species tree; b) species tree produced from BPP; c) *Haustorius allardi*; d) dorsal view of rostrum from *Haustorius galvezi*.

The parsimony tree produced from the character matrix generally recapitulated the results from BPP, with a few exceptions (**Fig. 3a**). BPP inferred *Neohaustorius* and *Acanthohaustorius* as being closer to the “*Haustorius-Lepidactylus*” clade than the character matrix supported, and that *L. dysticus* belonged to a clade consisting of *Parahaustorius* and *Protohaustorius*. This tree also differed somewhat from Sweet (1996), which inferred *Protohaustorius* as the most basal haustoriid.

### DIVERGENCE-TIME RESULTS

Dating analysis suggested that the haustoriids diverged from the gammaroids as early as the Paleocene (56–66 Mya; **Fig. 4**, **Fig. S5**). The subsequent split between the Pacific-endemic genus, *Eohaustorius*, and the remaining haustoriids followed shortly thereafter ~37 Mya (27–51 Mya, 95% HPD) during the Eocene. The East Asian *Eohaustorius* species were cut off from the North American continent by 25 Mya (16–37 Mya, 95% HPD). Further, we find evidence that the GoM was independently colonized by different genera of haustoriids at different times. The *Pseudohaustorius americanus* split is the deepest that includes a GoM endemic (~32 Mya), but without its Atlantic relatives we cannot conclude how long ago it colonized the Gulf. However, we do find that the ancestors of *Parahaustorius obliquus* split from its Atlantic-endemic relatives ~16 Mya, which makes it the earliest known colonization. The most abundant GoM endemics are those that appear to have colonized the Gulf most recently, within the last 10 million years: the *Acanthohaustorius*, *Haustorius*, and *L. triaticulatus* species groups. The most recent colonizations likely postdate the divergence of *H. canadensis* and *H. arenarius*, sister species separated on either side of the Atlantic (~9.5 Mya). Finally, we find that among the GoM-endemics, the deepest splits occur on opposite sides of the Mississippi River and likely occurred during the Pliocene or the end of the Miocene (2–7 Mya for *H. jayneae* and *H. galvezi*; 4–11 Mya for *L. triarticulatus* (GI-SR) and *L. triarticulatus* (PB-PC2)).

**Figure 4.**
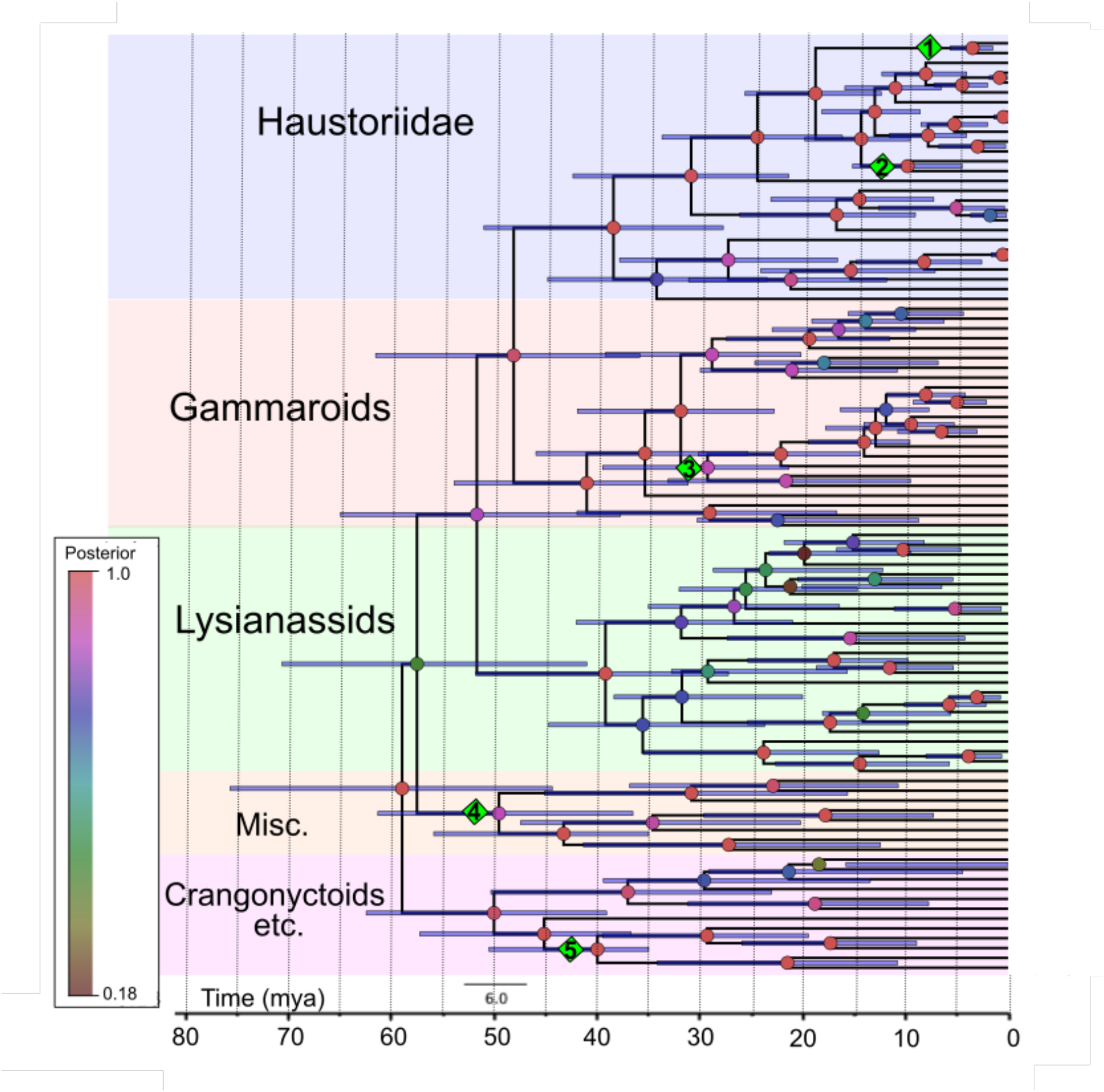
Time-calibrated phylogeny from BEAST2. Green diamonds represent calibration points: 1) closure of the Okefenokee Trough (1.75 Mya); 2) proposed migration to Europe by *H. arenarius* (~5 Mya); 3) Pontocaspian gammarid clade radiation (9–83 Mya); 4) Niphargidae-Pseudoniphargidae split (38–215 Mya); 5) Crangonyctidae-Pseudocrangonyctidae split (38–215 Mya).

### ANCESTRAL RECONSTRUCTION RESULTS

Ancestral range reconstructions indicated that haustoriids were likely originally subtidal species in the open ocean, and that the transition to the intertidal zone occurred twice: in *Eohaustorius* in the Pacific and in the *Haustorius-Lepidactylus* clade in the Atlantic (**Fig. 5**). Furthermore, we find that there were likely four independent shifts to brackish waters: *E. estuarius* in the eastern Pacific, *L. triarticulatus* complex in the GoM, *L. dysticus* in the Atlantic, and *H. allardi* in the Louisiana delta region (**Fig. 5**). In addition, the ancestral bottom preference was likely fine sand, with the Atlantic and eastern GoM endemics independently colonizing coarse sand (**Fig. S9**).

**Figure 5.**
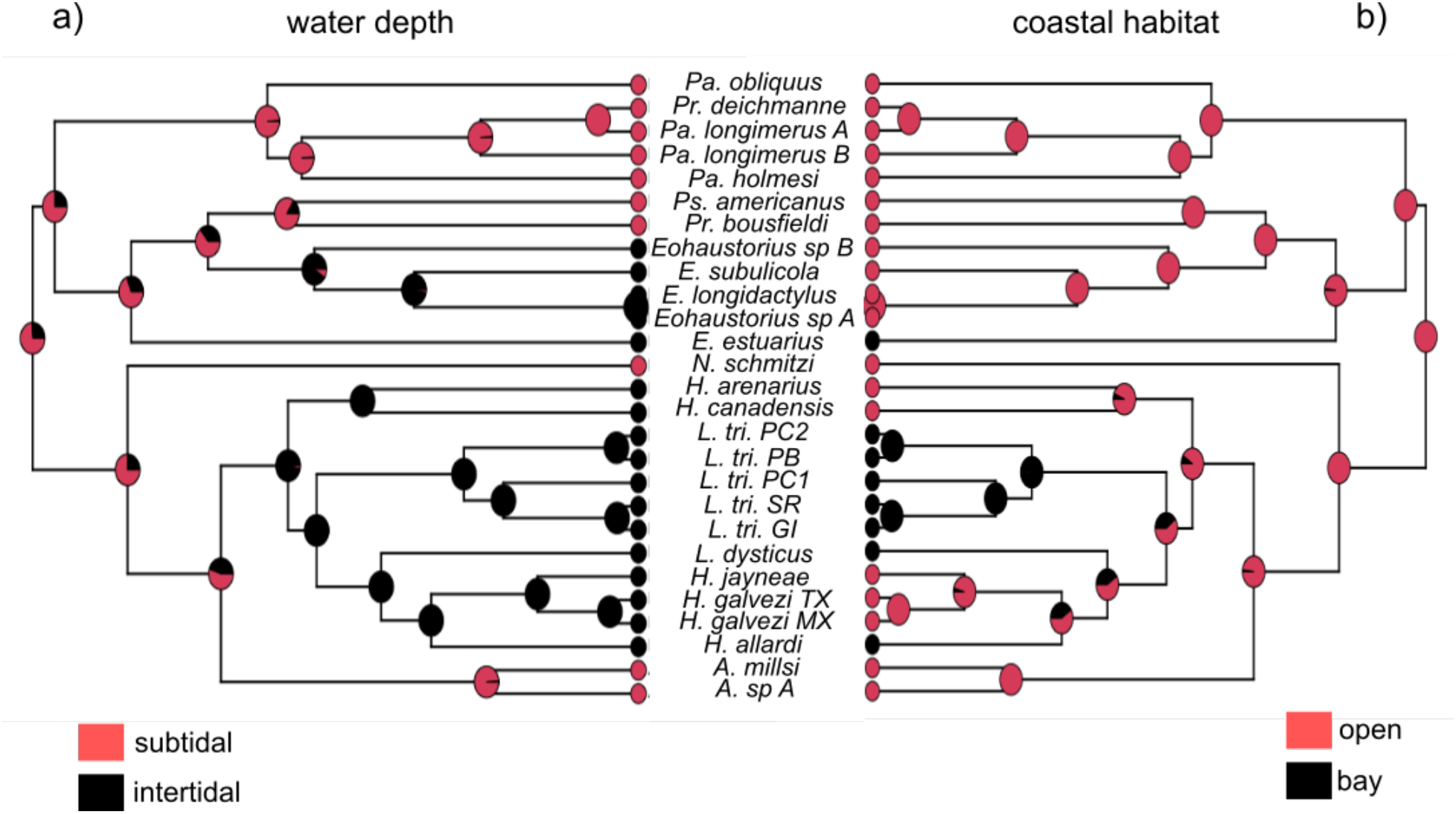
Ancestral habitat reconstructions. a) shifts between subtidal and intertidal; b) shifts between open and brackish coastlines. “L. tri” is short for *L. triarticulatus*, with initials following representing the lineage.

All reconstructed character traits were homoplastic to varying degrees (**Fig. S6–8**). Telson cleft and the number of spine groups on pereopod 7 article 4 were the least informative at the genus level (**Fig. S7b**, **S8a**). The size of the inner lobe of maxilla 2 relative to the outer was an inferred ancestral state for a *Pseudohaustorius-Eohaustorius-Protohaustorius* clade but may have arisen secondarily in *Neohaustorius* as well. The other two states of this character show little phylogenetic conservation and may not represent two distinct traits at all (i.e., they may be quantitative). The hooked state of epimeron 3 also supported this clade but there is a secondary loss of the hook in *Protohaustorius*.

The pleon character state (overhang or continuous) – a key trait in distinguishing *Lepidactylus* from *Haustorius* – has shifted from “overhang” (inferred to be ancestral) to “continuous” at least 4 times. Finally, the appearance of a distal lobe was inferred as derived in both *Eohaustorius* and GoM-endemic *Haustorius* species.

## DISCUSSION

In support of previous molecular phylogenies (Hou & Sket, 2016; Copilaş-Ciocianu *et al*, 2020), we find that haustoriid amphipods are basal gammarids and that there is no molecular support for the “Haustoriidira” parvorder (Lowry & Myers, 2017). Furthermore, we find that haustoriids in the Atlantic are monophyletic relative to those in the Pacific and likely diverged ~31 Mya. Finally, we find that the type-genus, *Haustorius*, is paraphyletic with respect to *Lepidactylus*. To resolve this issue, we recommend reserving *Haustorius* to Atlantic-specific species, placing the GoM-endemic “*Haustorius*” species within *Lepidactylus*, and elevating the GoM-endemic *L. triarticulatus* species complex to a new genus, *Cryptohaustorius*.

### HIGHER TAXONOMY OF AMPHIPODA

The higher taxonomy of Amphipoda has been so confused in the literature that Lowry & Myers (2017) noted that most guides present families in alphabetical order instead of phylogenetically. Several recent large-scale studies of amphipods have attempted to remedy this issue (Englisch *et al*., 2003; Havermans *et al*, 2010; Lowry & Myers, 2017; Hou & Sket, 2017; Copilaş-Ciocianu *et al*, 2020). In terms of the taxonomic placement of the Haustoriidae, a consensus has begun to emerge in molecular phylogenies of the group as basal gammarids (Englisch *et al*, 2003; Hou & Sket, 2017). These phylogenies have been contested by Lowry & Myers (2017) and Myers & Lowry (2018), who have argued that molecular phylogenies are “not built on any previous hypotheses” and are “not falsifiable”.

For our investigation of the higher taxonomy of amphipods, we have structured our analyses on existing competing hypotheses of Haustoriidira (Barnard & Drummond, 1982; Lowry & Myers, 2017). Specifically, we have examined the hypothesis that Haustoriidira is a monophyletic parvorder within the infraorder Lysianassida (see *Taxonomy* in **Methods** above). We find strong evidence that haustoriid amphipods are not related to the lysianassids but are instead basal gammarids as was proposed by Barnard & Drummond (1982). However, in opposition to Barnard & Drummond (1982) we find no support for the monophyly of the Haustorioidea; instead, the families generally allied in this group were found distributed across the amphipod phylogeny (**Fig. 2**). However, we have included far fewer members of the families outside Haustoriidae, and many families within the Haustoriidira are not represented in our phylogeny at all. Indeed, the vast majority have never been sequenced. Of special note is the lack of sequence data from the proposed sister to the Haustoriidae, the Condukiidae, and other Australian endemic families (i.e., Urohaustoriidae and the Zobrachoidae). Therefore, we can make no definitive statement as to the placement of these other families within the Amphipoda.

### BIOGEOGRAPHY OF THE HAUSTORIIDAE

Biogeographic hypotheses for the Haustoriidae were first proposed by Bousfield (1970), who argued that the family represented a recent adaptive radiation into an open beach habitat in the New England area and that they were actively displacing other fossorial amphipods along the North Atlantic coastline. Further, he posited that *Haustorius arenarius* diverged from *H. canadensis* as early as the Pleistocene during an interglacial period.

Our divergence-time analyses suggest a much earlier origin of the family (at least by the Eocene [37–47 Mya]; **Fig. 4**). Furthermore, we find evidence that the North Atlantic radiation began on the Eocene-Oligocene boundary (27–34 Mya). Interestingly, this corresponds to the Chesapeake Bay impact crater which decimated coastal marine life in the region and was followed by a period of rapid cooling (Ivany *et al*, 2000). This extinction event may have freed the coastal niche to new inhabitants, which was quickly exploited by the ancestors of the Haustoriidae. Furthermore, given that this event was followed by a period of cooling, it may help to explain the preference of haustoriid amphipods to cooler temperate climates. For example, haustoriid amphipods are largely absent from tropical areas in the Caribbean and South America despite the abundance of sandy beaches in these regions. Other explanations exist; for example, these niches may already have been filled by various species of mole crab (e.g. *Emerita portoricensis*) and isopods (e.g. *Exirolana* spp.), which are highly abundant in these regions (pers. obs.).

Haustoriids have nevertheless successfully colonized the warmer Gulf of Mexico on at least 4 separate occasions. The most abundant haustoriids are the intertidal species belonging to the genera *Lepidactylus* and *Haustorius*, which independently colonized the Gulf 6–15 Mya. Much of the Gulf states were submerged during this time as Pliocene sea levels were higher than they are today (Avise, 1992). The closing of the Isthmus of Panama during this epoch represented a dramatic alteration in the current regime as it forced the Loop Current over the submerged panhandle of modern Florida (Knowlton & Weigt, 1998). This increased oceanic current may have contributed to the divergence between Gulf and Atlantic species as has been proposed for many other coastal fauna (see Avise, 1992; Portnoy & Gold, 2012). Furthermore, multiple vicariant zones are known to exist within the GoM and have been proposed to explain divergence between sister taxa often separated by the Mississippi River or Mobile Bay (Portnoy & Gold, 2012; Hancock *et al*., 2019). These include the influx of freshwater down the Tennessee River ~2.4 Mya into Mobile Bay, the Suwannee Straits over northern Florida ~1.7 Mya, and increased sedimentation during the Miocene 5-10 Mya (Simpson, 1900; Bert, 1986; Portnoy & Gold, 2012; Hancock *et al*., 2019).

The Pacific-endemic haustoriid genus, *Eohaustorius*, diverged from the Atlantic group ~31 Mya (**Fig. 2**). The only species included in our phylogeny endemic to the western coastline of North America is *E. estuarius*, and it diverged from its western Pacific relatives ~25 Mya. In the western Pacific, *E. subulicola* was originally described from Japan (Hirayama, 1985), whereas *E. longidactylus* was first collected on the Korean peninsula (Jo, 1990). We collected both species on the Banzu tidal flats in Chiba, Japan. This could indicate that these species have historically been distributed across both Japan and the Korean peninsula; alternatively, the distribution could represent a secondary migration event after previously being separated by the Tsushima Straits (Takada *et al*, 2018). Indeed, these two species diverged ~2.5 Mya, which corresponds to the opening of the straits (Kitamura & Kimoto, 2006). During the Pleistocene (~10,000 years ago), a glacial maximum reconnected the Korean peninsula and Japan temporarily and could have facilitated secondary contact. On the other hand, in the Kisarazu region (including Banzu tidal flat) more than 1,800 tons of *Ruditapes philippinarum* (seed clams) are transplanted every year, but the source of the clams is not identified (Toba, 2002). Kitada *et al*. (2013) pointed out that large quantities of seed clams continue to be brought into Tokyo Bay from China. Okoshi (2004) reported that 22 species of marine benthic invertebrates were transported to Japan from Korean peninsula and Eastern China along with living clams. Therefore, further study is needed to determine whether *E. longidactylus* is native to Japan or was recently introduced by human shipping practices.

### TAXONOMIC RECOMMENDATIONS

We have shown that *Haustorius* is paraphyletic, and therefore suggest the following taxonomic rearrangements: 1) the genus name “*Haustorius*” is relegated to *H. canadensis* and the eastern Atlantic species; 2) GoM endemic “*Haustorius*” species represent a radiation of *L. dysticus* into the Gulf and therefore we suggest that these species should all be placed within the genus *Lepidactylus*; 3) the “*L. triarticulatus*” species complex is a distinct sister group to *Lepidactylus*, and we elevate this grouping to a new genus, *Cryptohaustorius*.

### GENUS *HAUSTORIUS* MÜLLER, 1775 WITH THE TYPE SPECIES *HAUSTORIUS ARENARIUS* SLABBER, 1769

#### Diagnosis

Large haustoriids restricted to open coasts in the Atlantic and Mediterranean. Maxilla 2 outer lobe is narrow and lanceolate, more than twice the size of the inner lobe. Pleon overhangs the urosome, with epimeron 3 smooth and ventrally curved (without hook). Telson cleft emarginate. Known species: *H. arenarius, H. canadensis, H. algeriensis, H. orientalis, H. mexicanus\** *The status of *H. mexicanus* is unclear. If a true *Haustorius* (in our definition above), it would represent the only *Haustorius* in the GoM. However, we have been unable to obtain specimens of this species and have not located it at its type locality.

### GENUS *LEPIDACTYLUS* SAY, 1818 WITH THE TYPE SPECIES *LEPIDACTYLUS DYSTICUS* SAY, 1818

#### Diagnosis

Small to medium size haustoriids in the GoM and Atlantic coastlines. May be found on open coasts or in brackish bays. Maxilla 2 outer lobe is narrow and lanceolate (except in *L. allardi*), and roughly twice the size of the inner lobe. Baler lobe on maxilliped is weakly developed. Pleon slightly overhangs the urosome, with epimeron 3 smooth and ventrally curved (without hook). Telson cleft emarginate or to base.

Known species: *L. dysticus, L. jayneae, L. galvezi, L. allardi*

### GENUS ***CRYPTOHAUSTORIUS GEN. NOV*.** WITH THE TYPE SPECIES *CRYPTOHAUSTORIUS TRIARTICULATUS* ROBERTSON & SHELTON 1980

#### Diagnosis

Small haustoriids (2–5 mm) restricted to the GoM. These are most numerous in brackish bays but may be found in small numbers subtidally on open coasts. Maxilla 2 outer lobe is broadly expanded and roughly twice the size of the inner lobe. Pleon is continuous with the urosome, with epimeron 3 smooth and ventrally curved (without hook). Telson cleft to the base, strongly spinated.

Etymology: From New Latin, the prefix “*crypto*” means “hidden” and the suffix “*haustorius*” is the type genus of the family.

Known species: *C. triarticulatus* (undescribed species complex; Hancock *et al*., 2019)

Type locality: Malaquite Beach, Texas

Known range: Northwestern Gulf of Mexico to at least Carrabelle Beach, Florida

Additional taxonomic issues may exist. For example, we found that the genus *Protohaustorius* appears to be polyphyletic, with *Pr. deichmanne* grouping within the *Parahaustorius* clade. However, without additional members of the genus and with low confidence of the phylogenetic position of *Pr. bousfieldi*, we suggest that additional data is needed before taxonomic reorganization is necessary.

## CONCLUSIONS

The Haustoriidae is a unique clade of sand-burrowing amphipods that has received little attention in recent decades. While considerable work remains to be done on the higher taxonomy of Amphipoda, it is now clear that the haustoriids are not relatives of the lysianassids. Instead, they represent an early split from gammaridean amphipods. Furthermore, other fossorial amphipods with similar morphology do not appear to be immediate relatives of the haustoriids, suggesting widespread convergence on traits ideal for a sand-dwelling lifestyle. Similarly, we find that most of the traits commonly used in dichotomous keys to distinguish the genera of haustoriids are homoplastic, suggesting need for systematic revision of taxonomically informative traits in Haustoriidae.

Future work should expand taxon sampling to the Australian-endemic “haustorioids”, such as the condukiids and the urohaustoriids, as well as include more representative members of other families within the “parvorder”. In addition, these phylogenies would benefit greatly from increased molecular resources including next-generation sequencing. A more exhaustive phylogeny of the fossorial amphipods may permit larger trait reconstructions to identify the timing of convergence to both phenotypes and habitat.

While the deconstruction of a taxonomic hypothesis in the absence of proposing a new higher classification scheme may seem like a step backwards, we contend that such a step is necessary to shed light on one of the many amphipod mysteries: the origin of the Haustoriidae.

## ACKNOWLEDGEMENTS

Special thanks to Gary Buhler and Aoi Tsuyuki for providing material for this study, and Denis Copilaş-Ciocianu of the Nature Research Centre in Lithuiana for sharing his alignments with us. Comments provided by Masanori of the Tokyo Bay Ecosystem Research Center has been a great help in this study. We are also particularly grateful to Dr. Hiroyuki Ariyama of Osaka Museum of Natural History and Yo Asada of Gifu University for advice on collection sites. We would also like to thank the many people who helped us collect in the field: Faith O. Hardin, Thomas Strawn, Shelby Strawn, Janelle Goeke and Justin Hilliard of Texas A&M University-Galveston, and Nuri Ubach, Sergio “Tama” Rodríguez, Jose “Pepe” Salgado, all of Unidad Académica Mazatlán, at the Instituto de Cinecias del Mar y Limnología, UNAM. This work was funded by grants from the Ecology & Evolutionary Biology (EEB) program at Texas A&M University, Texas Sea Grant, and Texas Ecolabs.

## SUPPLEMENTARY MATERIAL

Alignments and XML files are available at https://github.com/hancockzb.

### FIGURE CAPTIONS

**Figure S1.**
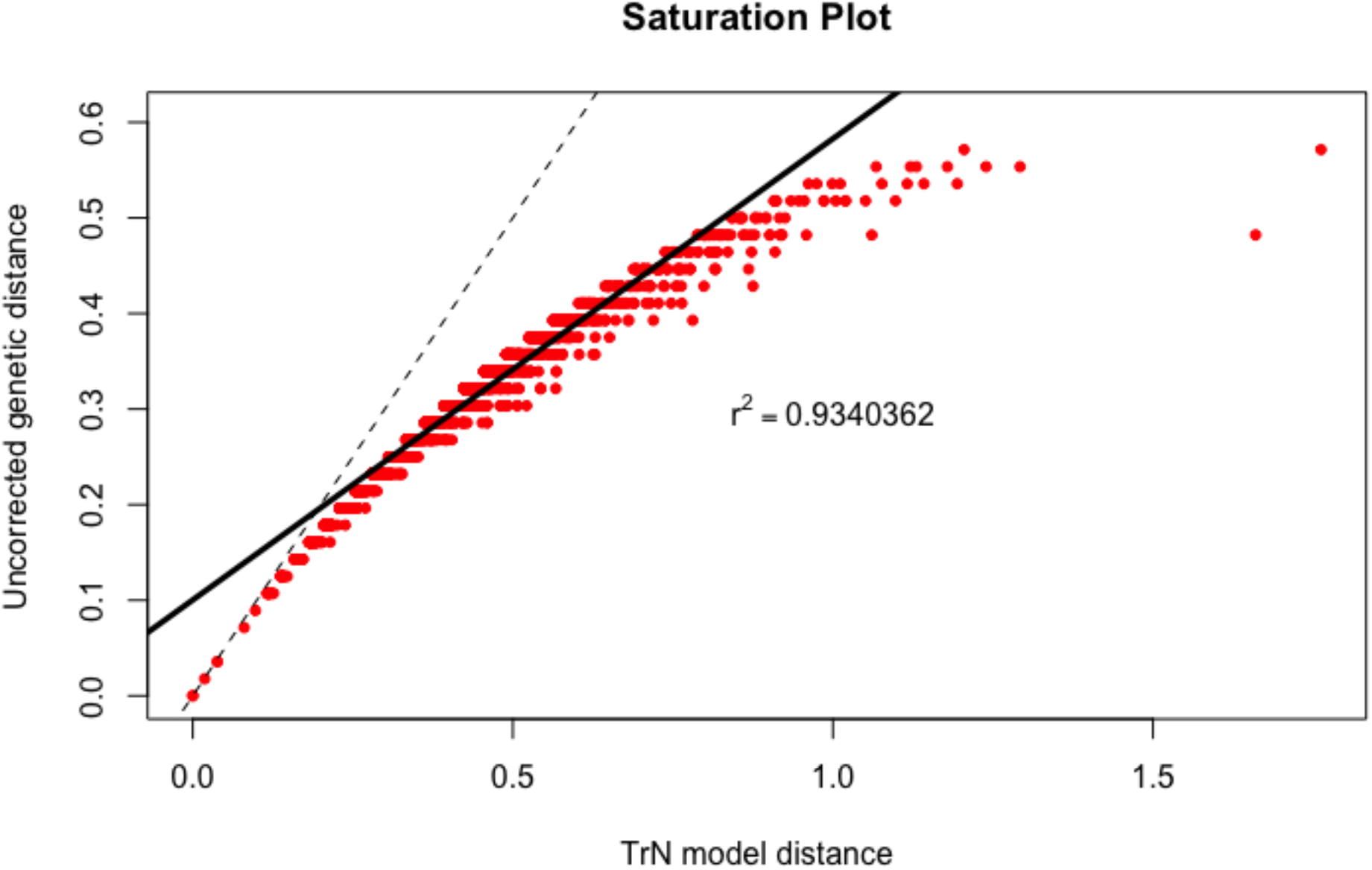
Saturation plot for *COI*.

**Figure S2.**
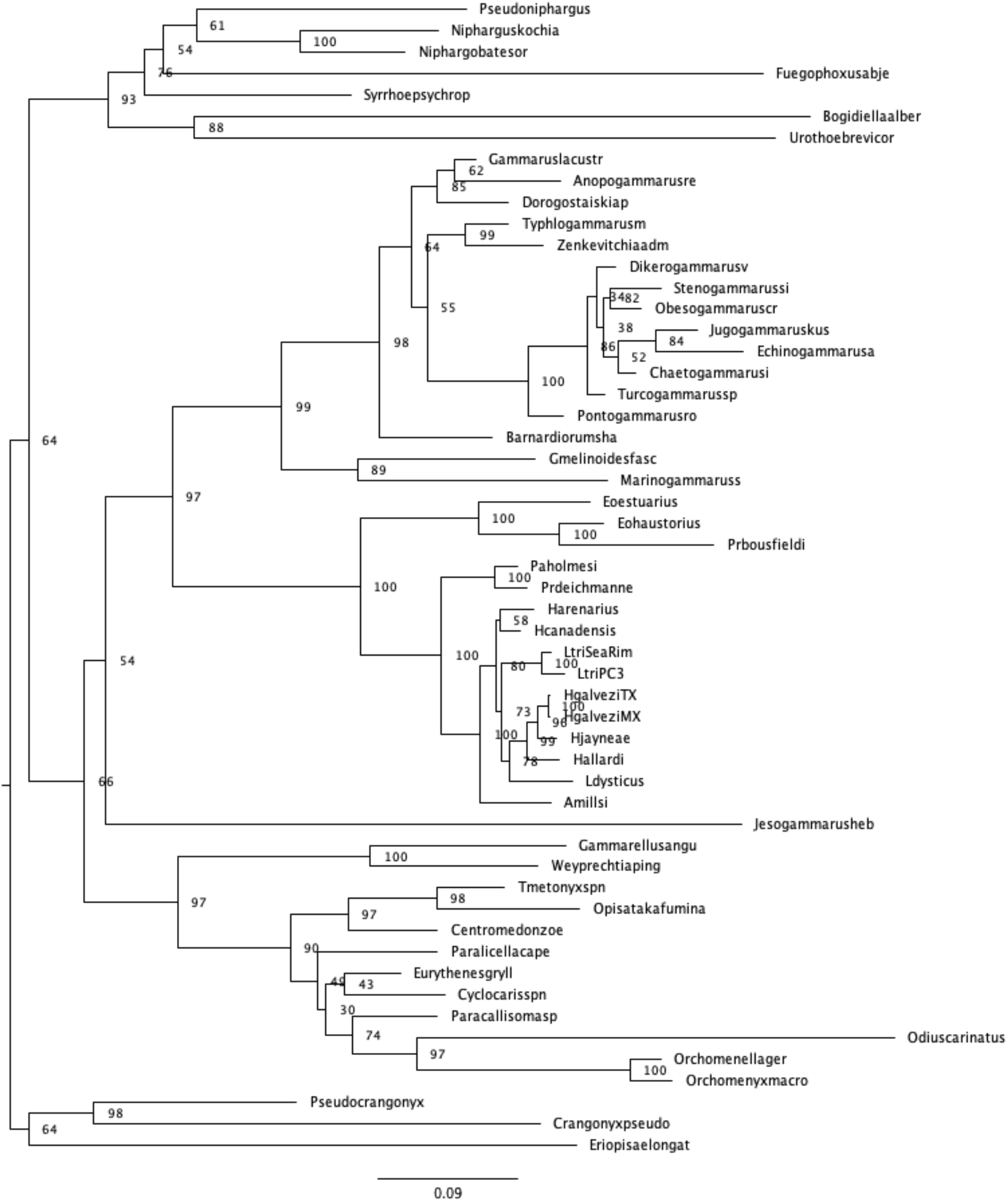
IQ-TREE phylogeny for the samples that passeMd the χ^2^ test. Node values represent aLRT scores.

**Figure S3.**
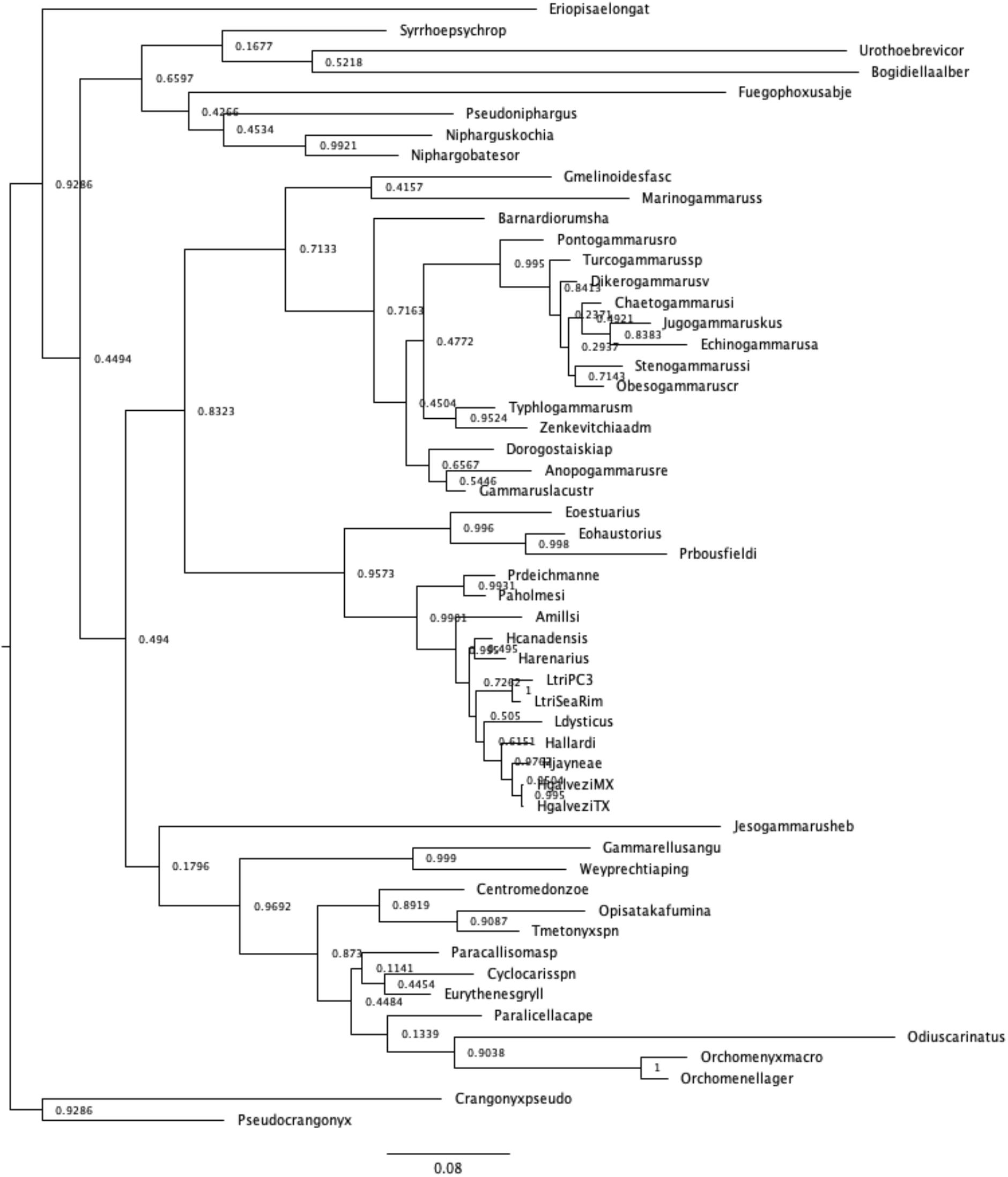
IQ-TREE phylogeny for the samples that passed the χ^2^ test. Bootstrap values shown at nodes.

**Figure S4.**
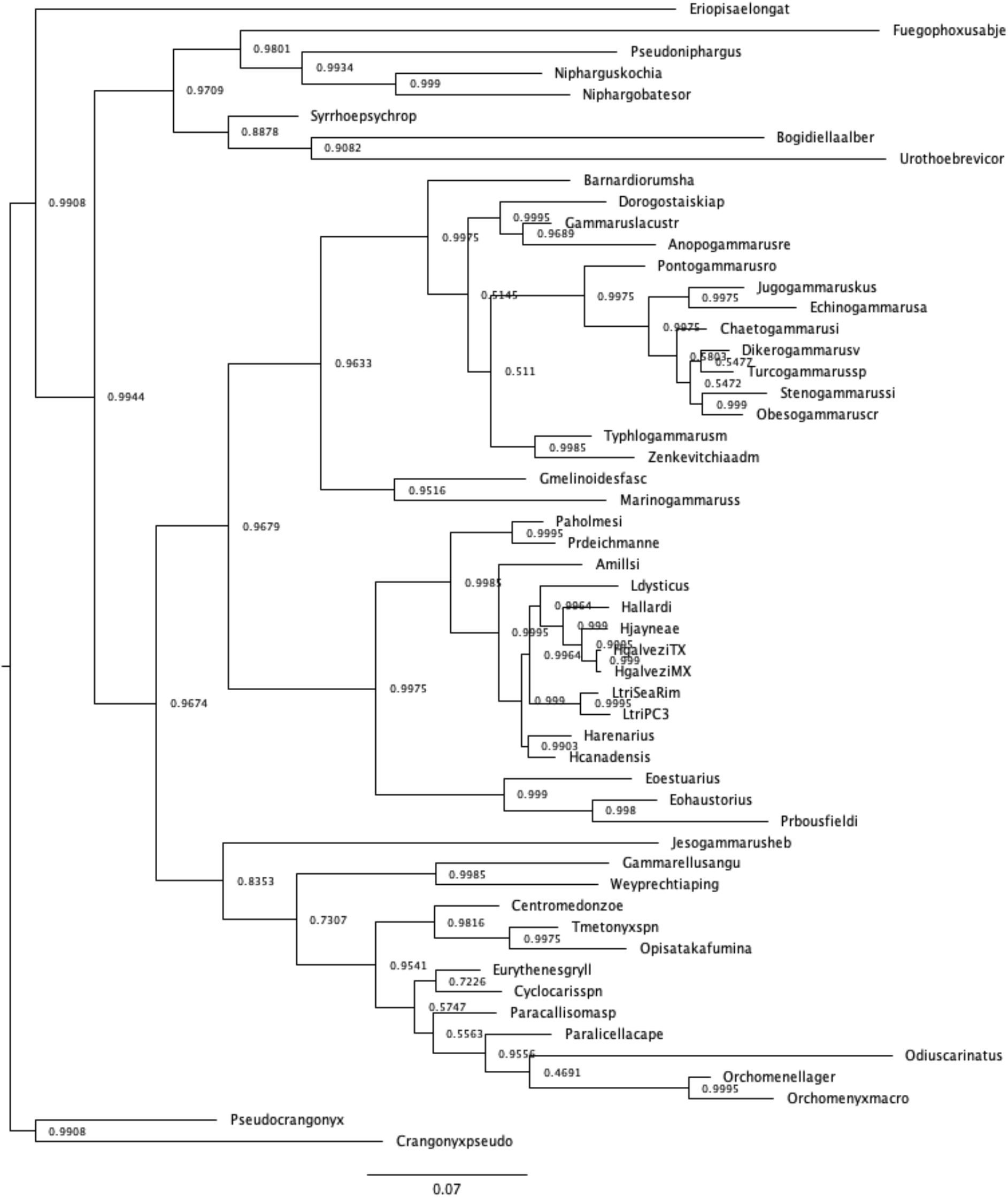
MrBayes phylogeny for the samples that passed the χ^2^ test. Posterior probabilities are shown at nodes.

**Figure S5.**
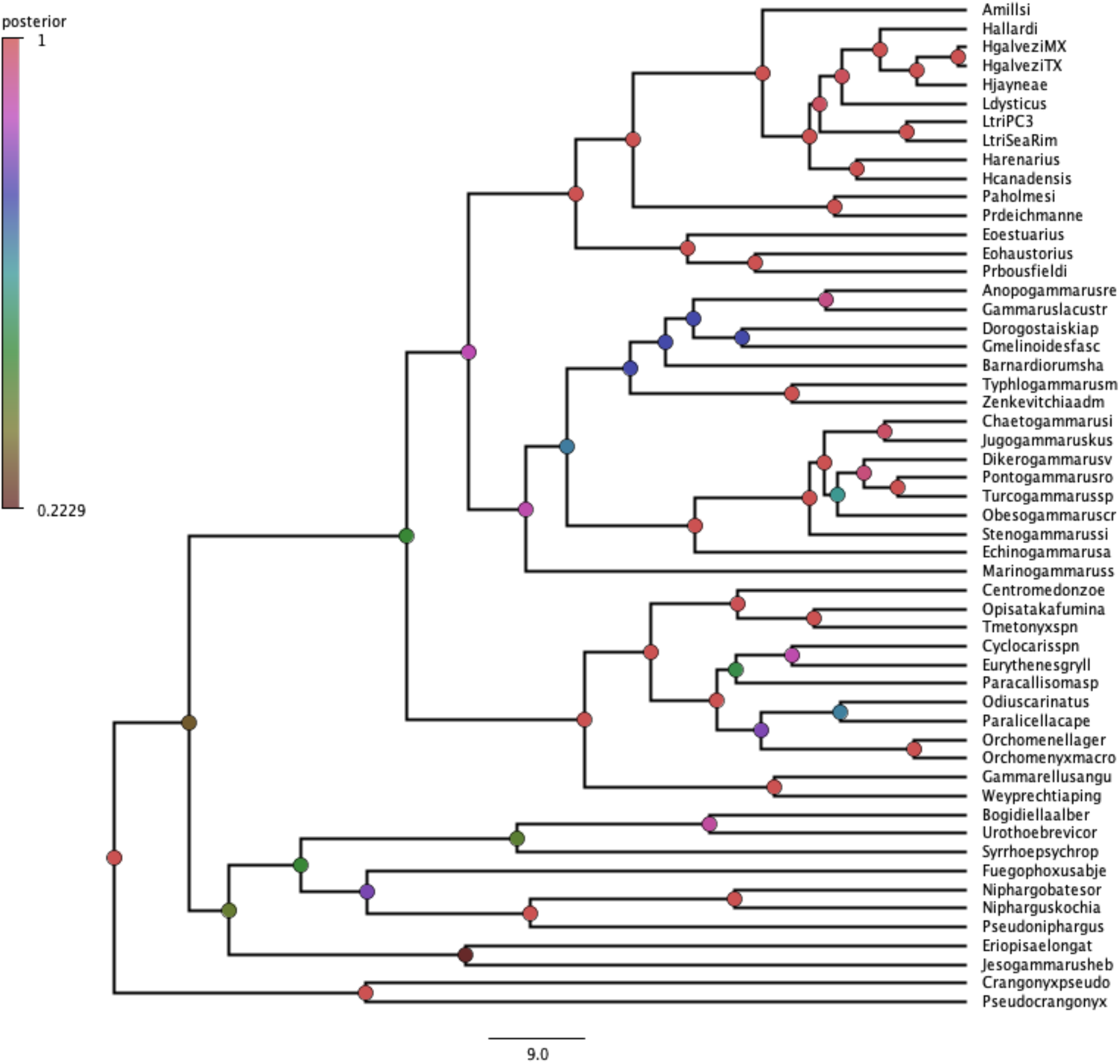
Calibration phylogeny from BEAST2 for the samples that passed the χ^2^ test.

**Figure S6.**
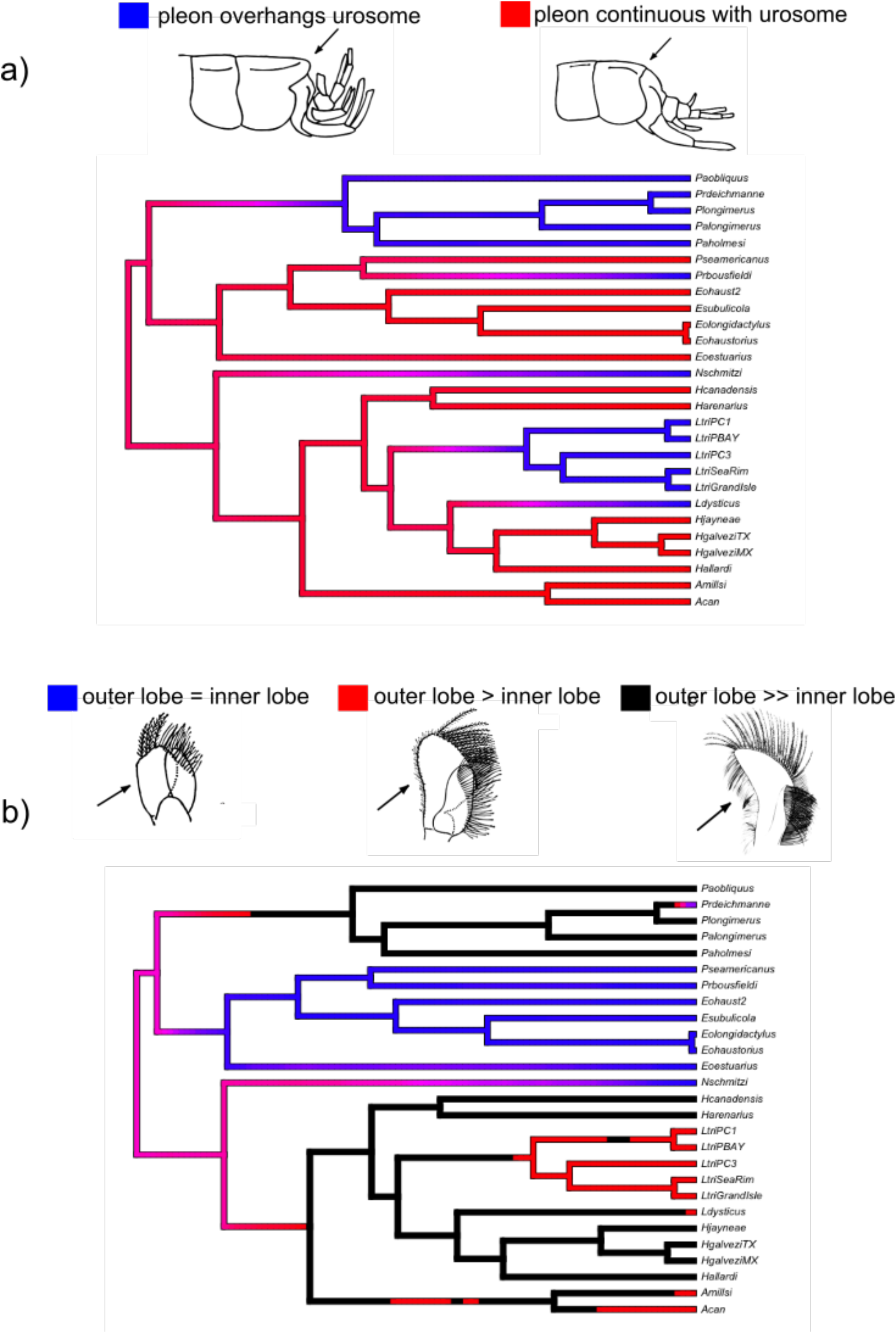
Ancestral reconstructions for morphological traits using *phytools*. Illustrations from LeCroy (2002) and Hancock & Wicksten (2018). A) Pleon overhangs urosome or not; B) maxilla 2 outer lobe size relative to inner lobe.

**Figure S7.**
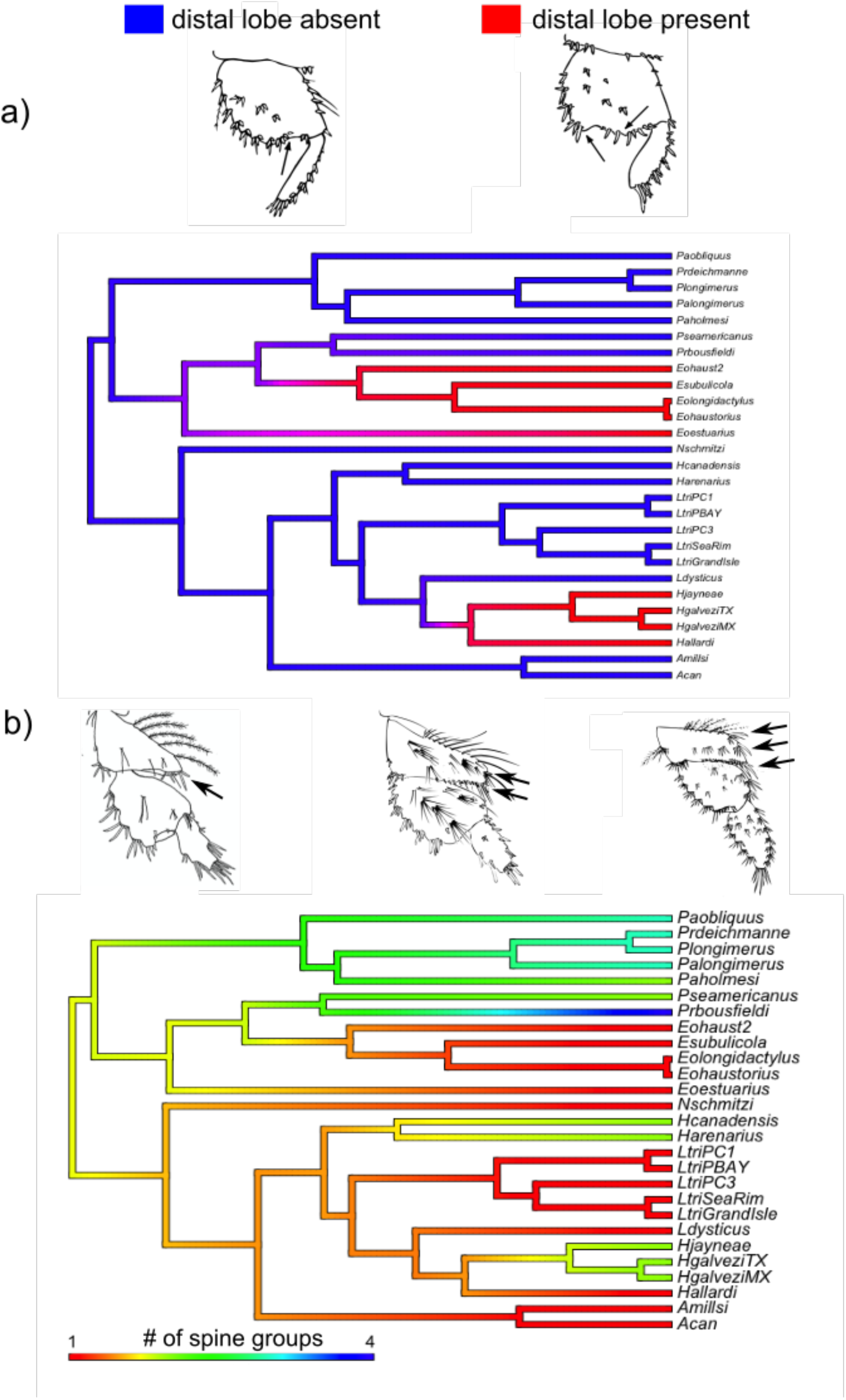
Ancestral reconstructions for morphological traits using *phytools*. Illustrations from LeCroy (2002) and Hancock & Wicksten (2018). A) Pereopod 6 article 5 distal lobe present or absent; B) pereopod 7 article 4 number of posterior spines.

**Figure S8.**
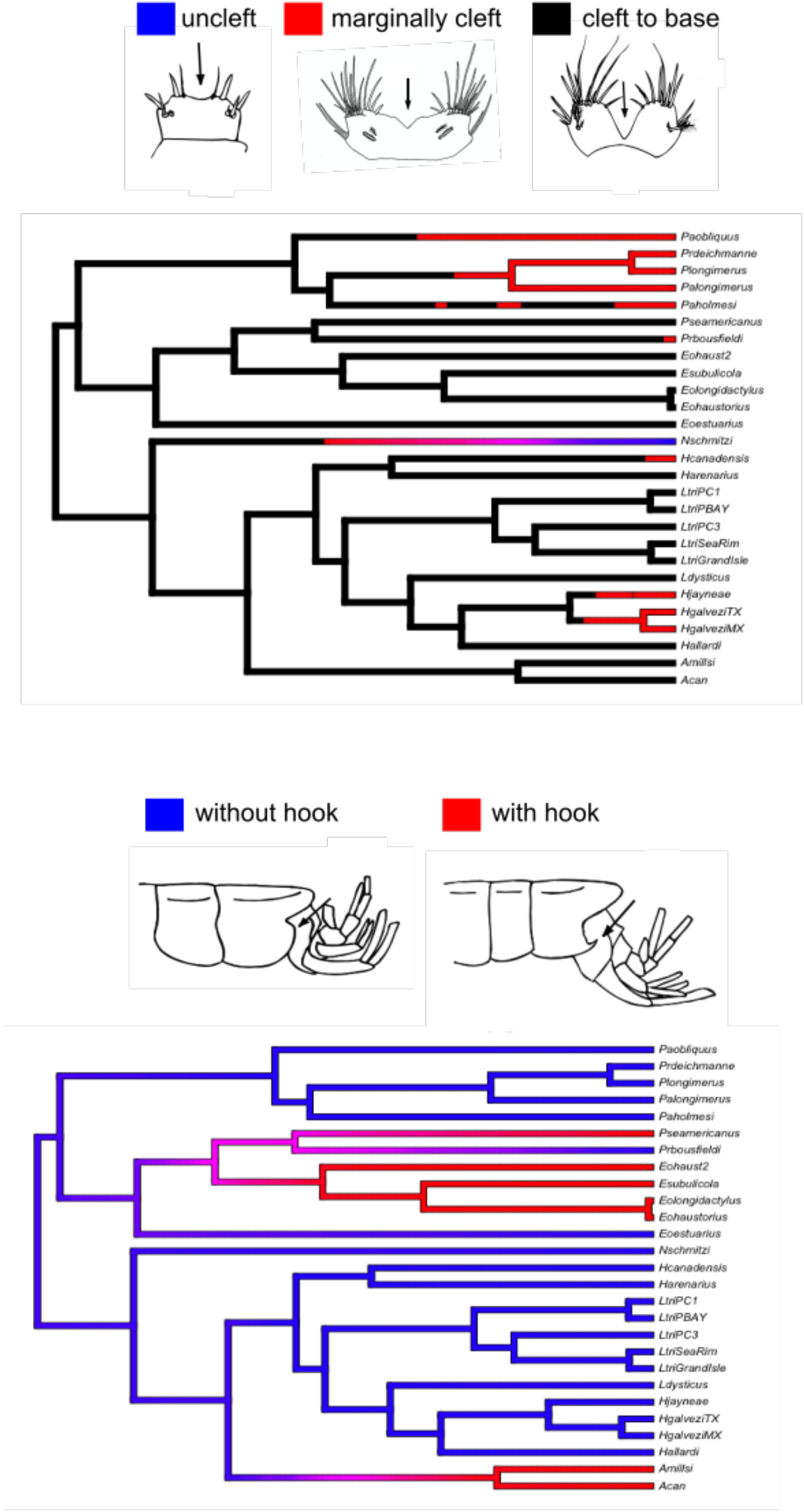
Ancestral reconstructions for morphological traits using *phytools*. Illustrations from LeCroy (2002) and Hancock & Wicksten (2018). A) Telsone cleft; B) epimeron 3 with or without hook.

**Figure S9.**
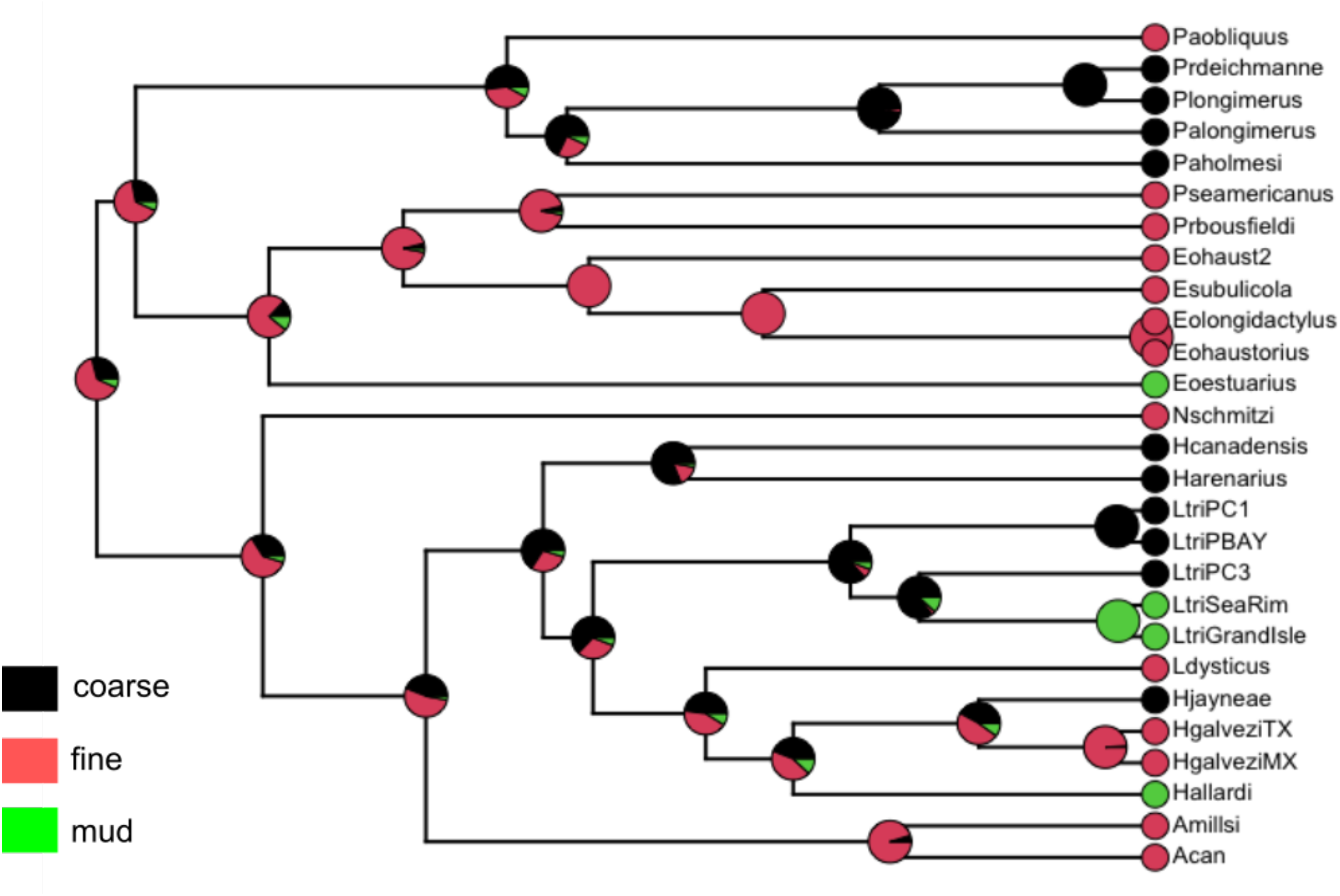
Ancestral reconstructions for bottom preference using *phytools*.

